# Ana1/CEP295 is an essential player in the centrosome maintenance program regulated by Polo kinase

**DOI:** 10.1101/2022.04.06.487296

**Authors:** Ana Pimenta-Marques, Tania Perestrelo, Patricia Rodrigues, Paulo Duarte, Mariana Lince-Faria, Mónica Bettencourt-Dias

**Affiliations:** Instituto Gulbenkian de Ciência, Rua da Quinta Grande, 2780-156 Oeiras, Portugal; CEDOC, Chronic Diseases Research Center, NOVA Medical School | Faculdade de Ciências Médicas, Universidade Nova de Lisboa, Lisbon, Portugal; Institute of Science and Technology Austria, 3400 Klosterneuburg, Austria

## Abstract

Centrioles play critical roles in our cells, being part of centrosomes and cilia, which are important microtubule organising centers (MTOC) with a variety of roles. While centrioles are very stable structures, they disappear in certain cell types upon differentiation, such as in oocytes. Little is known about the regulation of centriole structural integrity. We previously uncovered that the pericentriolar material (PCM), and its recruiter Polo kinase, are required for both the maintenance of centriole structural integrity and centrosome MTOC activity. Using an hypothesis driven RNAi screen, we show that both the cartwheel and the centriole wall play an important role in centrosome integrity. In particular, we uncovered that the centriole wall protein ANA1 is critical for the integrity of both new and mature centrioles, in *Drosophila* oogenesis as well as in cultured cells. Moreover, our results show that the activity of both Polo and the PCM in centriole integrity depends on ANA1. Our work suggests that the structural integrity of centrioles, once thought to be very stable organelles, depends on the turnover of key components, suggesting new perspectives for understanding the dysfunction of those structures in disease.

## Introduction

An important feature for cell homeostasis is how different structures are maintained in the cell. This is particularly relevant for organelles whose number and function are under tight control, as deregulation of these properties has critical implications for the cell. This is the case of the centrosome, the main microtubule-organizing center (MTOC) of eukaryotic cells. This organelle is composed of two centrioles, surrounded by a multiprotein matrix called the pericentriolar material (PCM) (Brito, Gouveia and Bettencourt-Dias, 2012; Conduit, Wainman and Raff, 2015). The PCM is indispensable for centriole biogenesis and for nucleating and anchoring MTs at the centrosome (Pimenta-Marques and Bettencourt-Dias, 2020). Centrioles are small microtubule (MT) cylinders with a striking 9-fold radial symmetry of doublets or triplets of MTs that build up the centriole wall **(Fig. 1B)**. At their most proximal part, centrioles have a cartwheel structure consisting of a central hub and nine radially-arranged spokes along their length (Callaini, Whitfield and Riparbelli, 1997; Guichard, Hamel and Gönczy, 2018). The cartwheel is composed of the conserved proteins SAS6 and ANA2/STILL (Kitagawa *et al*., 2011; Dzhindzhev *et al*., 2014; Cottee *et al*., 2015). Proteins such as BLD10/CEP135, ANA1/CEP295 and SAS4/CPAP localize more externally, at the centriolar wall (Fu and Glover, 2012; Mennella *et al*., 2012; Sonnen *et al*., 2012; Tian *et al*., 2021). BLD10/CEP135 and ANA1/CEP295 interact (Fu *et al*., 2016) and extend from the inner to the most outer part of the centriole, where the C-terminus of ANA1 is positioned between the MT blades (Tian *et al*., 2021). Both ANA1 and SAS4 interact with different PCM proteins (Gopalakrishnan *et al*., 2011; Conduit *et al*., 2015; Fu *et al*., 2016; Galletta *et al*., 2016). At the distal tip of the centriole a cap is found which is composed of CP110 and CEP97 (Kleylein-Sohn *et al*., 2007). While clear appendages have been found at the distal end of the centriole in many other organisms, this is not the case in Drosophila cultured cells, which form no cilia and do not have strong astral microtubule arrays (Franz *et al*., 2013; Gottardo, Callaini and Riparbelli, 2015). The number of centrioles, and consequently the number of centrosomes, is tightly controlled in actively dividing cells. Centrosome dysfunction is associated with a variety of human diseases including cancer (Cirillo, Gotta and Meraldi, 2017; Goundiam and Basto, 2021) and microcephaly (Jayaraman, Bae and Walsh, 2018). Historically, centrosomes have been regarded as exceptionally stable structures. They are resistant to drug- and cold-induced depolymerization (Kochanski and Borisy, 1990) and to forces and MT destabilisation at the entrance of mitosis (Belmont *et al*., 1990). Furthermore, experiments in *C. elegans* showed that the pool of the centrosomal proteins SAS-6 and SAS-4, present in the centrioles and contributed by the sperm, persist for several embryonic cell cycles with no detectable exchange with the cytoplasmic pool (Balestra, Von Tobel and Gönczy, 2015), which suggests that centrioles are stably inherited through many divisions.

**Figure 1-.**
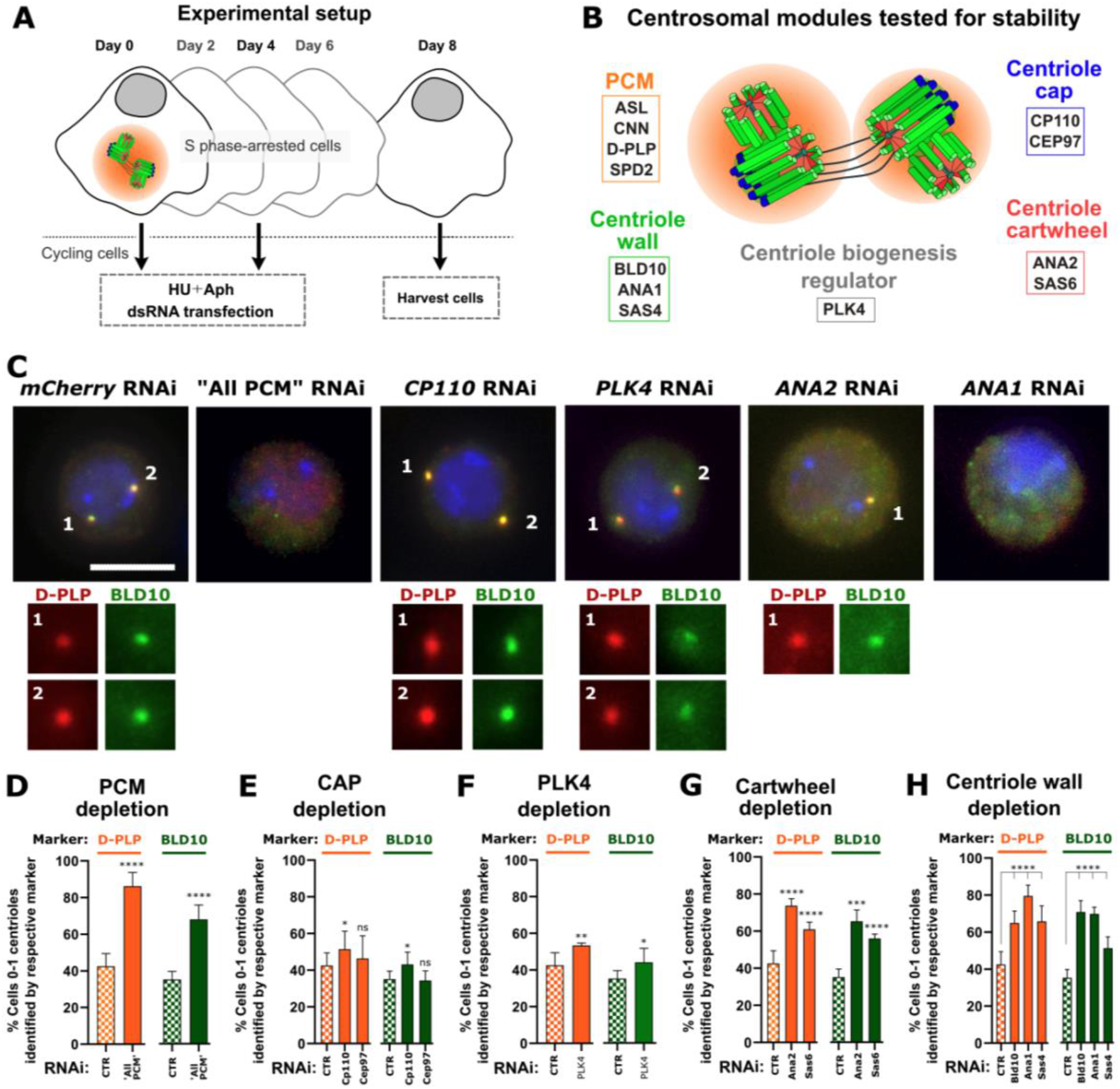
The PCM, the cartwheel and the centriole wall are critical for the maintenance of centriole structural integrity. **(A)** Schematic representation of the “centriole stability assay”: cells were transfected at day 0 with double stranded RNA (dsRNA), and simultaneously arrested in S-phase with HU (hydroxyurea) and Aph (aphidicolin). On day 4, cells were subjected to a second round of dsRNA transfection and treatment with HU and Aph. On day 8, cells were harvested and assayed for centriole numbers by immunofluorescence. **(B)** Schematic representation of molecules depleted in the screen. **(C)** Representative images of centrosomes stained with D-PLP (red) and BLD10 (green) in control cells (mcherry RNAi) and cells with dsRNA transfection for “All PCM” (ASL, CNN, D-PLP and SPD2 as a positive control as previous described in), PLK4, CP110, ANA2, or ANA1. Scale bar, 5 µm. DNA (blue). Enlargements of centrosomes present in each cell are shown. **(D-H)** Centriolar numbers were assessed considering the staining for the PCM marker D-PLP (orange bars) or centriole wall marker BLD10 (green bars). Quantification of the percentage of cells with abnormally low centriole numbers (0-1 centrioles) upon depletion of **(D)** “All PCM”, **(E)** CP110 and CEP97, **(F)** PLK4 kinase, **(G)** SAS6 and ANA2, and **(H)** BLD10, ANA1 and SAS4. Data are the mean ± SEM of three independent experiments (n ≥ 100 cells per condition in each experiment). Generalised linear binomial model in which the effect of each replicate and RNAi were considered as independent factors. * *p*< 0.05; ** *p* < 0.01; \*\*\**p* < 0.001; \*\*\*\**p* < 0.0001; ns, not significant. (see also **Fig. S1**).

Despite their inherent stability, centrosomes are lost from the oocytes of most metazoan species (Manandhar, Schatten and Sutovsky, 2005; Werner, Pimenta-Marques and Bettencourt-Dias, 2017) and are known to be inactivated (i.e. loss of their MTOC capacity) in some cell types that undergo differentiation, such as neuronal, muscle and epithelial cells (Muroyama and Lechler, 2017; Sanchez and Feldman, 2017). Several studies in different proliferating cell types have uncovered the pathways regulating how centrosomes mature and become active MTOCs. However, far less is known on the pathways regulating centrosome inactivation or elimination in different cell types. These are important questions to understand how abnormal loss or abnormal maintenance of centrosomes in development or disease might affect cell homeostasis.

We have previously identified what we named a *centrosome maintenance mechanism*, which operates in the *Drosophila* germ-line and somatic cells (Pimenta-Marques *et al*., 2016). This process is led by Polo (PLK1 in Humans), a conserved kinase that drives PCM recruitment (Lane and Nigg, 1996; Dobbelaere *et al*., 2008; Haren, Stearns and Lüders, 2009) and PCM maintenance (Singh, Ramdas Nair and Cabernard, 2014; Pimenta-Marques *et al*., 2016; Cabral *et al*., 2019). This mechanism is shut down in the female germ-line, with loss of Polo and the PCM from centrosomes, followed by centrosome functional inactivation and loss of centrioles, demonstrating that these players are also critical for centriole structural integrity (Pimenta-Marques *et al*., 2016). The depletion of four major PCM proteins (ASL, SPD-2, CNN, and D-PLP) or of Polo in S-phase arrested *Drosophila* cultured cells, leads to centriole number reduction (Pimenta-Marques *et al*., 2016), suggesting that this maintenance mechanism also plays an important role in other cells. These findings point to the importance of the recruitment of Polo and PCM components to the centrosome, supporting the existence of a regulated homeostatic maintenance program that is switched off in acentrosomal cells (Pimenta-Marques et al., 2016). How Polo and the PCM promote centriole structural integrity is not known. It is possible that those molecules prevent the loss of essential centriole components. Indeed, there is some evidence supporting the dynamicity of PLK1/PCM/centriole components (Keller *et al*., 2014; Novak *et al*., 2014; Woodruff, Wueseke and Hyman, 2014; Conduit *et al*., 2015).

Here we first test whether different centrosome components have a role in centriole structural integrity by conducting a RNAi screen targeting components of different centrosome structural parts. We show that the cartwheel and the centriolar wall are critical for centriole structural integrity. We further identify an essential role for the centriolar wall protein ANA1 in centriole integrity, with its removal leading to the disappearance of fully matured centrosomes. Finally, we found that both Polo and the PCM require ANA1 to promote centriole structural integrity.

## RESULTS

### The centrosome is a dynamic structure, with the PCM, the centriole wall and the cartwheel being critical for its integrity

Previous evidence shows that despite centriole structural intrinsic stability, some of its components are dynamic (Bahmanyar *et al*., 2010; Mahjoub, Xie and Stearns, 2010; Conduit *et al*., 2014, 2015; Keller *et al*., 2014; Woodruff, Wueseke and Hyman, 2014), suggesting they might be essential for centriole stability, together with the PCM and Polo (Pimenta-Marques *et al*., 2016). To test this hypothesis, we conducted a screen to test whether different centriole components can have a role in centrosome integrity.

We used a “centriole stability assay” that we previously developed and validated (Pimenta-Marques *et al*., 2016) **(Fig. 1 A)**. In this assay, *Drosophila* tissue culture cells (DMEL) are arrested in S-phase, which halts the centriole biogenesis cycle after centriole duplication (Dzhindzhev *et al*., 2010). This procedure maintains the number of centrosomes constant, which allows the uncoupling of the maintenance of centrosome integrity from centrosome biogenesis. We targeted different parts of the centrosome using RNAi, which should only affect centriole structure if those components are involved in stability and need to be replenished constantly (as RNAi prevents the translation of new proteins). Such components include proteins of different centriole substructures such as: the centriole core, the cartwheel; the centriole wall, and the centriole cap. We also targeted the PCM as a positive control, given its role in the centrosome maintenance program (Pimenta-Marques *et al*., 2016). We depleted proteins known to be important for these different modules **(Fig. 1 B, Fig. S 1)**: **1)** the building blocks of the cartwheel: SAS6 (Kitagawa *et al*., 2011; Cottee *et al*., 2015) and ANA2/STIL (Dzhindzhev *et al*., 2014; Cottee *et al*., 2015); **2)** the centriolar wall: BLD10/CEP135 (Kleylein-Sohn *et al*., 2007; Roque *et al*., 2012), SAS4/CPAP (Kleylein-Sohn *et al*., 2007; Gopalakrishnan *et al*., 2011), and ANA1/CEP295 (Chang *et al*., 2016; Fu *et al*., 2016); **3)** the cap at the distal end of centrioles: CP110 and CEP97 (Kleylein-Sohn *et al*., 2007; Fu and Glover, 2012); and **4)** the PCM: for this module, 4 major PCM proteins were simultaneously depleted (Martinez-Campos *et al*., 2004; Dzhindzhev *et al*., 2010; Fu and Glover, 2012; Mennella *et al*., 2012, 2014; Fu *et al*., 2016), as individual depletion was previously shown not to be sufficient to induce centriole loss (ASL, CNN, D-PLP and SPD2 (“All PCM)) (Pimenta-Marques *et al*., 2016). We also tested PLK4 kinase as it is a major regulator of centriole biogenesis, known to regulate several centriolar components (Bettencourt-Dias *et al*., 2004, 2005; Habedanck *et al*., 2005; Zitouni *et al*., 2014). To infer which parts of the centrosome were disturbed upon RNAi, we used markers of the PCM (D-PLP), the centriolar wall (BLD10, ANA1, and SAS4), and the distal centriole cap (CP110) **(Fig. 1 B-H**; and **Fig. S 1)**.

As previously shown by us (Pimenta-Marques *et al*., 2016), simultaneous depletion of the four major PCM proteins (ASL, CNN, D-PLP and SPD2 – “All PCM”) induced a strong reduction of centriole number, confirming the dynamicity and importance of the PCM for the maintenance of centrosome integrity **(Fig. 1 C,D**; and **Fig. S 1 B)**.

Cells depleted of cap proteins, despite showing a reduction in SAS4 foci numbers **(Fig. S 1 C)**, did not show a strong reduction in the other markers **(Fig. 1 C,E**; and **Fig. S 1 C)**. It is possible that loss of SAS4 upon cap protein depletion reflects a specific interaction with CP110 (Galletta *et al*., 2016). Similarly, to the cap proteins, PLK4 depletion did not lead to centriole loss, judged by the different markers analysed **(Fig. 1 C,F, Fig. S 1 D)**. Nonetheless, we observed loss of SAS4 foci **(Fig. S 1 D)**, suggesting that besides the known role of PLK4 in the recruitment of CPAP (the Human counterpart of SAS4) to centrioles (Moyer and Holland, 2019), PLK4 might also be involved in SAS4 maintenance at *Drosophila* centrioles. In contrast to its essential role in centrosome biogenesis, our data shows that PLK4 is not critical for the maintenance of centrosome integrity, reinforcing that centriole biogenesis and maintenance pathways are differentially regulated. This result is in line with other experiments where Drosophila or Human cells were subjected to PLK4 inhibition for a very long time and centriole number was reduced as a consequence of loss of duplication, but loss of centriole integrity was not observed (Wong *et al*., 2015; Nabais *et al*., 2021).

On the other hand, depletion of cartwheel or centriolar wall proteins led to a strong decrease in centriole number, measured by the presence of the PCM (PLP) and centriolar proteins BLD10, SAS4, ANA1, and CP110 **(Fig. 1 C,G,H** and **Fig. S 1 E,F)**. These results suggest that individual proteins from both the wall and the cartwheel are dynamic and critical for the maintenance of the whole centrosomal structure. Moreover, it is likely that maintenance of both assembling daughters, as well as mother centrioles, requires the recruitment of proteins that compose those substructures, as most cells analysed had only 0 or 1 centrioles.

Together our results suggest that the PCM, the wall, and the cartwheel require a turnover of components which contribute to centrosome maintenance, in particular centriole structural integrity. From all the candidates tested, depletion of ANA1 led to the strongest effect on different centrosomal markers **(Fig. 1 C,H Fig. S 1 F)**, similar to “All PCM” depletion **(Fig. 1 C,D Fig. S 1 B)**. Interestingly, ANA1/CEP295 has been shown to function as a centriolar bridge, connecting the centriole wall with the PCM (Fu *et al*., 2016; Tsuchiya *et al*., 2016; Tian *et al*., 2021). The direct interaction between ANA1 and the PCM protein ASL in Drosophila (Fu *et al*., 2016) and with Cep152 (Fu *et al*., 2016) and Cep192 in Humans (Tsuchiya *et al*., 2016) is required for centriole-to-centrosome conversion (Izquierdo *et al*., 2014; Fu *et al*., 2016). This process is important for the stability of centrioles after the cartwheel loss observed at the exit of mitosis in human cells (Izquierdo *et al*., 2014). In addition, phosphorylation of ANA1 at S-S/T motifs was recently shown to prompt the recruitment of Polo to mother centrioles, contributing to PCM maturation and centriole elongation (Alvarez-Rodrigo *et al*., 2021). Also, In *Drosophila*, ANA1 is one of the last proteins to be lost from centrioles before they are eliminated in oogenesis (Pimenta-Marques *et al*., 2016), and from the ommatidia of the eye (Riparbelli *et al*., 2018). Similarly, it is one the last proteins to be lost upon centrosome reduction in the sperm basal body (Blachon *et al*., 2009; Khire *et al*., 2016). Altogether these evidences suggest that ANA1 may be an important player in the PCM/Polo centrosome maintenance mechanism. Therefore, we further investigated the role of ANA1 in centrosome maintenance, in particular centriole structural integrity, in fully matured centrosomes, exploring its function *in vivo*, as well as its mechanism of action in relation to Polo and the PCM.

### The centriolar wall protein ANA1 is required for centrosome maintenance *in vivo*

The female germline is a great system to study the maintenance of centrosomes as these structures are progressively eliminated throughout the different stages of oogenesis. In early stages, oocytes are specified from cysts of 16 interconnected cells. The centrosomes from 15 cells migrate into the oocyte forming a large MTOC which is active up to mid stages of oogenesis (stages 6-7). At these stages, centrosomes start first losing Polo and the PCM, followed by their progressive elimination before meiotic metaphase I (Pimenta-Marques *et al*., 2016). Therefore, we investigated if ANA1 is important for centrosome maintenance *in vivo* in this system.

We depleted a large portion of existing ANA1 using the deGradFP system (**Fig. 2**). This system can target endogenous GFP-tagged proteins for degradation by the proteasome through tissue-specific expression of a modified F-box protein that is fused to an anti-GFP nanobody (Caussinus, Kanca and Affolter, 2012). The deGradFP system is independent of protein turnover, relying on the ubiquitin-proteasome pathway to degrade GFP tagged proteins. We specifically induced the expression of deGradFP in oocytes from stages 3/4 onward, when most centrosomes should have already duplicated and migrated from the nurse cells to the oocyte (Mahowald and Strassheim, 1970). Here we expressed ANA1-GFP under its endogenous promoter in the genetic background of *ANA1* mutant (Blachon *et al*., 2008). Centrioles were analysed at stages 10 of oogenesis, where they are still present in control conditions (**Fig. 2 A**) (Pimenta-Marques *et al*., 2016). Expression of deGradFP led to approximately 80% decrease in the total levels of ANA1-GFP on centrioles (Degron condition), when compared to ANA1-GFP flies without deGradFP expression (control, **Fig. 2 B**), showing that a significant pool of ANA1 is being depleted. Given that centrioles are densely packed at late stages of oogenesis, we measured the total intensity of different markers as a proxy for centriole content as previously done (Pimenta-Marques *et al*., 2016). We used D-PLP and γ-tubulin as PCM markers, and CP110 as a centriole marker, as they are robust and convenient markers in these cells. ANA1 depletion led to a strong reduction in all markers, suggesting centrioles are being prematurely lost **(Fig. 2 B)**. Collectively, our observations show that ANA1 is important for maintaining the structural integrity of centrioles, and consequently to maintain fully mature centrosomes in different cell types.

**Figure 2-.**
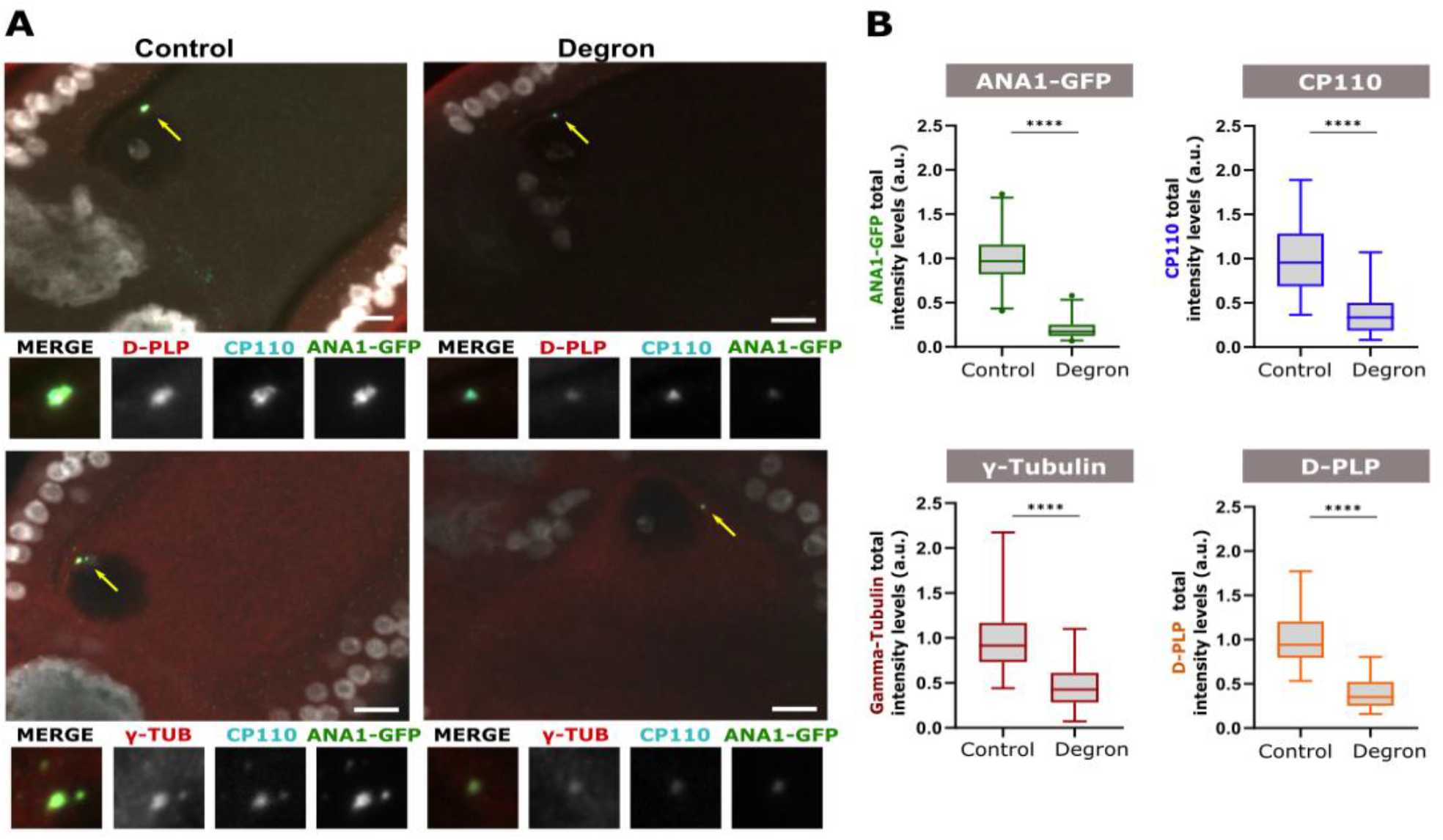
The centriolar wall protein ANA1 is required for centrosome maintenance *in vivo*. **(A-B)** Expression of the deGradFP tissue-specific system in oogenesis (using UAS/Gal4) leads to tissue-specific degradation of ANA1-GFP (expressed under the control of its endogenous promoter). deGradFP (Degron Condition) was induced in the germ line after stages 3/4 onward, a stage where centrosomes have duplicated and migrated to the oocyte. ANA1-GFP flies without the deGradFP system were used as controls. **(A)** Stage 10 oocytes were immunostained for centriole marker CP110 (cyan), D-PLP (red) and γ-TUB (red). (**B)** Quantification of total intensity levels of ANA1-GFP, CP110, D-PLP and γ-Tub. The decrease of ANA1-GFP levels in oogenesis leads to a strong reduction in the levels of PCM (γ-TUB and PLP) and centriole (CP110) markers analysed. More than 25 oocytes were analysed per condition. Enlargements of the indicated areas are shown. Scale bars, 10 µm. Box-and-whisker plot (whiskers extend to the 2.5th and 97.5th percentiles) of the total integrated intensity. \*\*\*\**p* < 0.0001 (unpaired Mann-Whitney test).

### ANA1 is a player in the Polo mediated centrosome maintenance program

We have previously found that Polo kinase and the PCM are critical for the maintenance of centrosome integrity. Tethering Polo to centrosomes using the PACT domain of the PCM protein D-PLP (Gillingham and Munro, 2000; Martinez-Campos *et al*., 2004) rescues the loss of PCM and the loss of centrosomes, both in culture cells depleted of PCM, and in oogenesis where the PCM is naturally lost (Pimenta-Marques *et al*., 2016). Whether ANA1 could play a role in the Polo-mediated centrosome maintenance program is not known. Therefore, we investigated whether ANA1 and Polo synergise in the centrosome maintenance pathway.

We first asked whether Polo requires ANA1 to prevent centriole loss. We used the “centriole stability assay” in *Drosophila* culture cells, where we depleted the PCM to trigger centriole loss **(Fig. 3 A)**. Expression of GFP-PACT was used as a control. As expected, cells depleted of “All PCM” and expressing GFP-PACT (control) have abnormally low numbers of centrioles (0-1 centrioles) (**Fig. 3 B** and **Fig. S 2 A,B)**. In this context, as previously shown, expression of GFP-Polo-PACT partially rescues centriole loss, when compared to cells expressing GFP-PACT only (**Fig. 3 B** and **Fig. S 2 A,B)** (Pimenta-Marques *et al*., 2016). However, upon “All PCM” and ANA1 depletion, GFP-Polo-PACT expression was no longer capable of rescuing centriole numbers (**Fig. 3 B** and **Fig. S 2 A,B**). These results show that the partial rescue provided by GFP-Polo-PACT is dependent on ANA1.

**Figure 3-.**
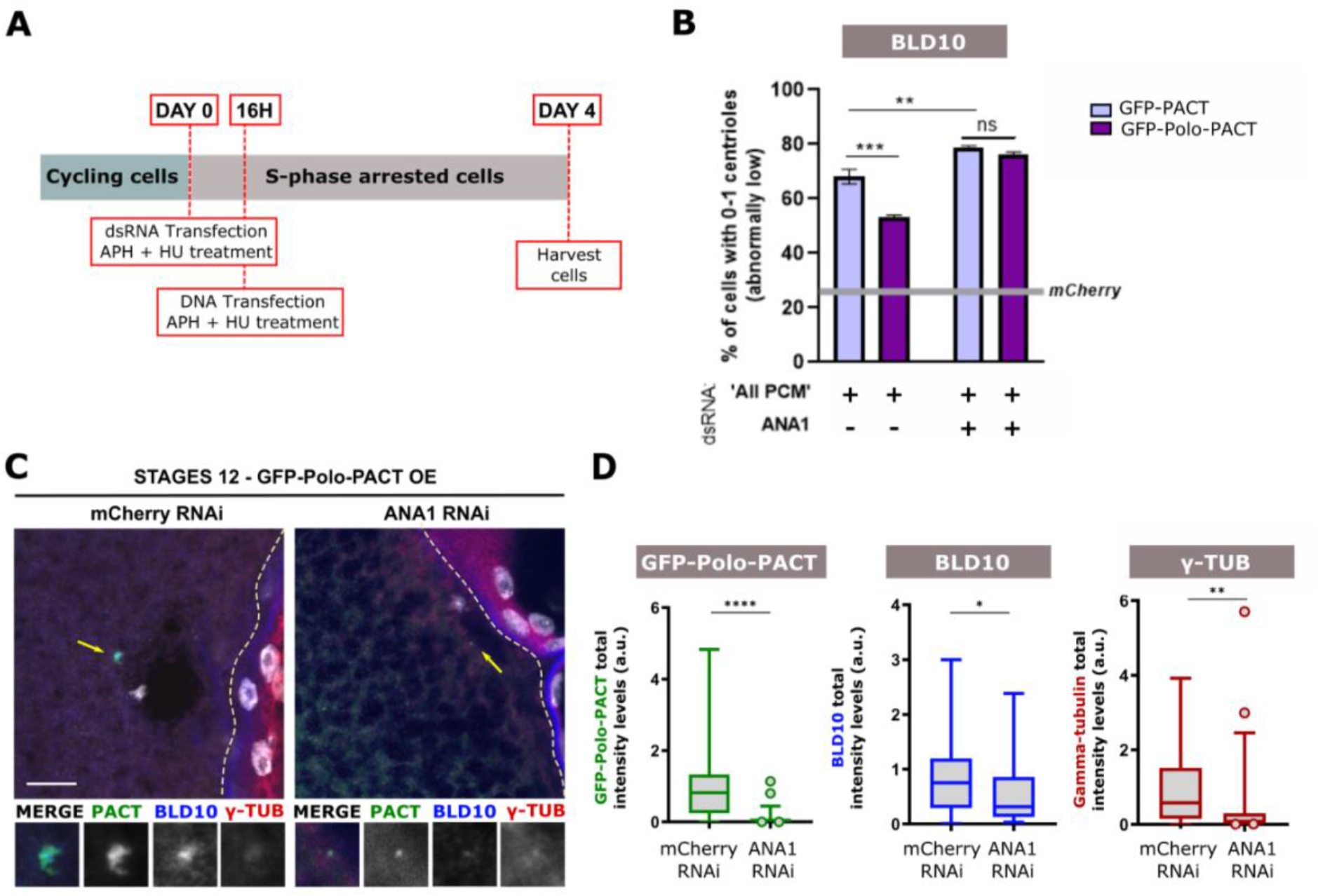
Centrosome maintenance by Polo kinase is dependent on the centriolar wall protein ANA1. **(A)** Schematic representation of the experimental setup: DMEL cells were subjected to dsRNA transfection and treatment with Aph (aphidicolin) and HU (hydroxyurea) at day 0. Cells were depleted of “All PCM” as before, or simultaneously depleted of “All PCM*”* and ANA1. After 16h, cells were transfected (GFP-PACT or GFP-Polo-PACT) in medium with Aph and HU. Cells were harvested and assayed for centriole numbers by immunofluorescence at day 4. *mCherry* RNAi was used as a negative control. **(B)** Quantification of the percentage of cells with abnormally low numbers of centrioles (i.e. 0-1). Centrioles were identified by considering the positive staining in each cell for the centriolar wall protein BLD10. The grey line represents the percentage of cells with 0-1 centrioles in the control (cells transfected with *mCherry* dsRNA and expressing GFP-PACT). Note that simultaneous depletion of “All PCM’’ and ANA1 had a stronger effect on centriole loss when compared to depleting the “All PCM” alone. Bars represent the mean ± SEM of three independent experiments (n>100 cells per condition in each experiment), Two-way ANOVA, with Tukey’s multiple comparisons test (**C)** Stage 12 egg chambers expressing GFP-Polo-PACT (centrosome maintenance conditions) and ANA1 RNAi or mCherry (control for RNAi). Enlargements of the indicated areas (yellow arrows) are shown. Scale bars, 10 μm. (**D)** Quantification of the total intensity levels of different centrosomal proteins (GFP-Polo-PACT, BLD10 and γ-TUB) in stages 12 of oogenesis, in GFP-Polo-PACT expressing oocytes. Box-and-whisker plot (2.5th and 97.5th percentiles) of the total integrated intensities of the different markers analysed. A minimum of 28 oocytes were quantified for each condition, unpaired Mann-Whitney test. For all the statistical tests used in this figure: *, *p*<0.05; ** *p*< 0.01; *** *p*<0.001; \*\*\*\**p*<0.0001; ns, not statistically significant.

In oogenesis, tethering Polo to the oocyte centrioles leads to a significant maintenance and/or recruitment of γ-tubulin to these structures. Under these conditions, centrioles are maintained up to meiotic metaphase I, a stage where they are naturally absent. These centrioles contain ANA1 and are active MTOCs (Pimenta-Marques *et al*., 2016). We thus asked if ANA1 is required for Polo-induced centriole maintenance *in vivo*. GFP-Polo-PACT was expressed after stages 3/4 of oogenesis while ANA1 synthesis was prevented by RNAi (**Fig. 3 C,D)**. In stages 12 of oogenesis, when the majority of centrioles are normally lost in control conditions (GFP-PACT expressing oocytes), as previously described, tethering Polo to the centrioles led to maintenance of centrosomes, as seen by γ-tubulin and BLD10 **(Fig. 3 C,D; Fig. S 2 C)**. However, upon ANA1 RNAi, the levels of both γ-tubulin and BLD10 were significantly reduced, and there was a clear reduction of the percentage of stage 12 egg chambers showing the presence of centrioles **(Fig. 3 C,D** and **Fig. S 2 C)**. Therefore, our results suggest that, as observed in cultured cells, Polo-induced centrosome maintenance is dependent on ANA1.

### Ana1 is sufficient for maintaining centriole structural integrity in oogenesis

A scenario compatible with our results is that ANA1 is a critical structural component of the centrioles that is constantly replenished by new protein translation, and that Polo and the PCM are important to ensure ANA1’s role at the centrioles. If that is the case, then ensuring the incorporation of ANA1 at the centrioles should be sufficient to ensure their structural integrity, even when the PCM and Polo are low. We thus tested if tethering ANA1 to the oocyte centrioles is sufficient to maintain them until later stages (stages 12), when most egg chambers have already lost their centrioles.

To tether ANA1 and other proteins to the centriole, we used a more generalised strategy. We developed a nanobody trapping experiment, using the anti-GFP single domain antibody fragment (vhhGFP4) (Saerens *et al*., 2005; Caussinus, Kanca and Affolter, 2012) fused to the PACT domain of D-PLP (Gillingham and Munro, 2000) to predominantly trap GFP tagged proteins to the oocytes centrioles. By analysing egg chambers in stages 10 of oogenesis, a stage where centrioles are still normally present, we show that the PACT::vhhGFP4 is efficient for tethering GFP-tagged proteins to the centrioles **(Fig. 4 A,F)**. As a positive control, by using this system, GFP-Polo increased γ-tubulin levels on the oocyte centrioles in comparison with the control (tethering of GFP alone) **(Fig. 4 A,C)**, as previously observed for GFP-Polo-PACT (Pimenta-Marques *et al*., 2016). The levels of ANA1 at the oocytes centrioles in stages 10 were also significantly increased by expressing ANA1-GFP with the PACT::vhhGFP4 **(Fig. 4 A,B)**, showing that ANA1 is efficiently tethered at centrioles. Interestingly, the tethering of ANA1 to the centriole increases mildly the levels of BLD10, suggesting that ANA1 may contribute to stabilise the centriole structure by maintaining the levels of other important wall proteins **(Fig. 4 A,D)**. However, in contrast to tethering of Polo, ANA1 did not promote additional recruitment/maintenance of the PCM component, γ-tubulin **(Fig. 4 A,C)**.

**Figure 4-.**
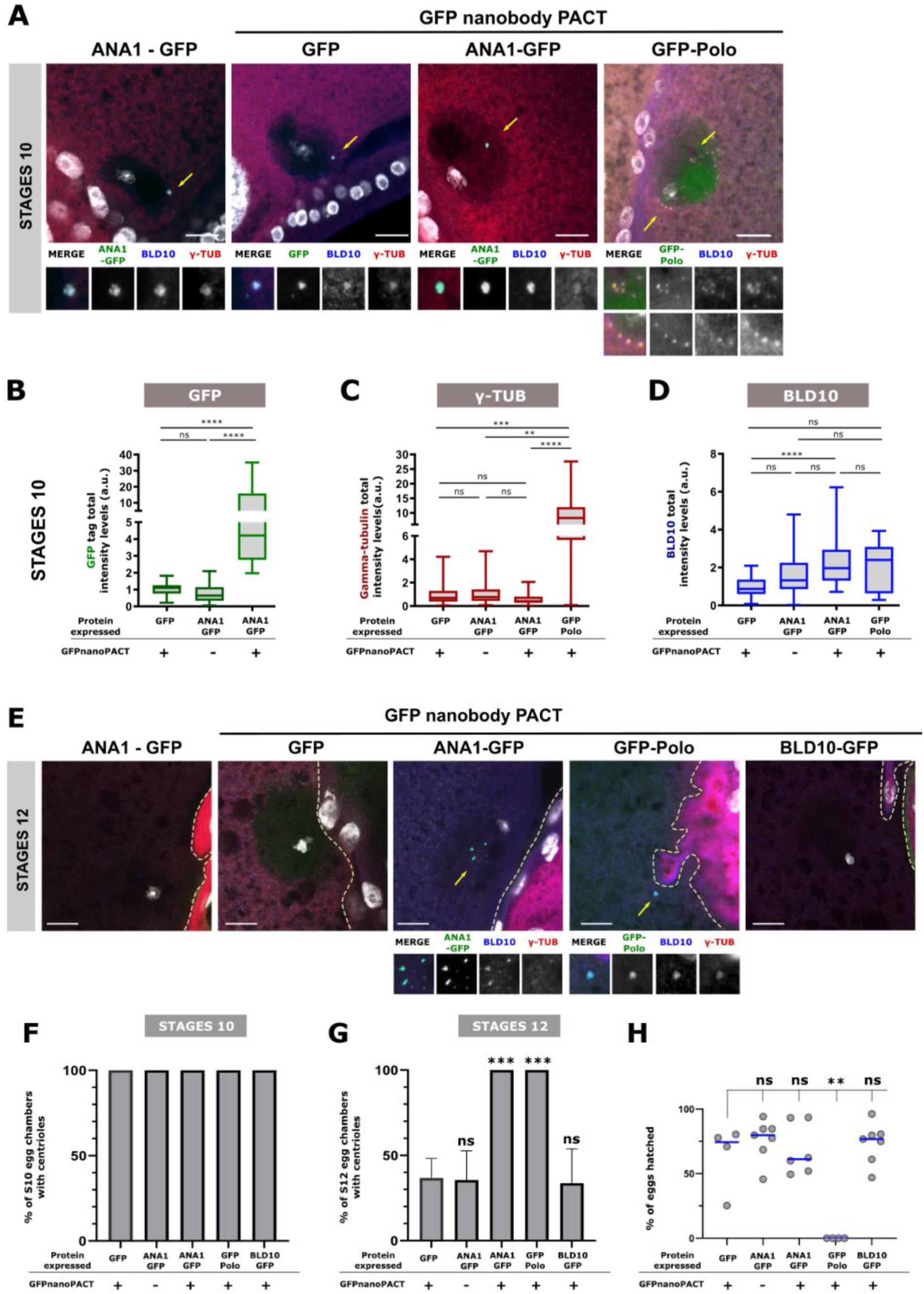
Ectopic tethering of ANA1 to the oocyte centrioles extends centriole presence up to stages 12 of oogenesis. **(A-D)** Analysis of stage 10 oocytes upon tethering different centrosomal proteins to the oocyte centrioles by expressing a GFP nanobody construct fused to the PACT domain (PACT::vhhGFP4) that targets molecules to the centriole. Enlargements of the indicated areas (with centrioles, yellow arrows) are shown. Note that at this stage the oocyte is supposed to have approximately 64 clustered centrioles, known to be scattered when Polo-PACT is expressed (Pimenta-Marques et al., 2019). Scale bars, 10 μm. **(B)** Quantification of the total intensities of GFP and ANA1-GFP tethered to centrioles by PACT::vhhGFP4, as well as ANA1-GFP without tethering to the oocyte centrioles (without PACT::vhhGFP4 expression) in stages 10. A minimum of 28 oocytes were analysed for each condition. **(C-D)** Quantification of the total intensities of **(C)** γ-tubulin and **(D)** BLD10 in stage 10 egg chambers expressing either GFP, GFP-Polo or ANA1-GFP in combination with PACT::vhhGFP4. Quantifications were also performed for stage 10 egg chambers expressing ANA1-GFP without PACT::vhhGFP4 expression. Note that tethering GFP-Polo leads to an increase in the total levels of γ-tubulin on the oocyte centrioles in stages 10, which is not observed upon forced localization of ANA1 on the oocyte centrioles. A minimum of 27 oocytes were analysed for each condition, with the exception of GFP-Polo condition in which a total of 15 oocytes were analysed. Box-and-whisker plot (whiskers extend to the 2.5th and 97.5th percentiles). **(E)** Representative images of each tested condition. Enlargements of the indicated areas (yellow arrows) are shown. Scale bars, 10 μm. Percentage of egg chambers from **(F)** stages 10 and **(G)** stages 12 showing the presence of centrioles (centrioles identified by the colocalization of the GFP signal with Bld10 staining). Between 26 and 30 oocytes were quantified for each condition, with exception of GFP-Polo expression where n=14. Statistical significance was determined by Fisher’s Exact test on the pooled replicates, after testing whether there were significant differences between them. P-values were Bonferroni corrected for a Family Wise Error Rate of 5%. Shown are the mean ± SEM. **(H)** Quantification of the number of eggs hatched from the total number of eggs laid by females expressing GFP; GFP-Polo; ANA1-GFP and BLD10-GFP in the presence of PACT::vhhGFP4 and females expressing ANA1-GFP alone without PACT::vhhGFP4. Each dot in the plot represents the percentage of eggs hatched from the total number of eggs laid by a single female. (Kruskal-Wallis test). For all the statistical tests used in this figure: *, *p*<0.05; ** *p*< 0.01; *** *p*<0.001; \*\*\*\**p*<0.0001; ns, not statistically significant.

We then analysed stages 12 of oogenesis, when centrioles normally start to be eliminated and asked whether ANA1 would be sufficient to keep their structural integrity, even if not capable of retaining their PCM. This experiment is particularly important to understand whether ANA1 is indeed a critical player in centriole integrity, downstream of the PCM. In the control condition, where GFP was tethered to the centriole, only ∼30% of stage 12 oocytes showed the presence of centrioles (colocalization of GFP with BLD10) **(Fig. 4 E,G)**. As expected, GFP-Polo/PACT::vhhGFP4 expressing flies showed 100% of stage 12 oocytes with the presence of centrioles **(Fig. 4 E,G)**. When tethering ANA1 to the oocyte centrioles (ANA1-GFP/PACT::vhhGFP4 expressing oocytes), 100% of stage 12 oocytes showed the presence of centrioles **(Fig. 4 E,G)**. This data demonstrates that ANA1 is capable of maintaining centrioles in a scenario where both Polo and PCM are naturally down-regulated (Xiang *et al*., 2007; Jambor *et al*., 2015; Pimenta-Marques *et al*., 2016). Importantly, this is likely to be a phenotype specific to ANA1, as tethering another centriolar wall protein, BLD10, was not sufficient to prevent normal centriole elimination **(Fig. 4 E,G)**. Our data shows that ANA1 is critical for maintaining centriole integrity, independently of a role in PCM recruitment. Interestingly, tethering of ANA1 to centrioles did not lead to any obvious defects in meiosis, in contrast to tethering of Polo (data not shown, (Pimenta-Marques *et al*., 2016). Moreover, flies expressing ANA1-GFP/PACT::vhhGFP4 were fertile and laid eggs which hatched at a comparable rate as control flies (expression of GFP/PACT::vhhGFP4) **(Fig. 4 H)**. These centrioles which were structurally maintained are most likely inactive as they do not recruit PCM **(Fig. 4 C)** and do not nucleate MTs, which could interfere with the meiotic spindle and subsequent embryonic nuclear divisions, as observed upon Polo tethering (Pimenta-Marques *et al*., 2016). Our observations *in vivo* suggest that maintaining a given threshold of ANA1 at the centriole provides stability to this structure, even when the PCM levels are low. All together, these data show that ANA1 is a critical player in providing integrity to centrioles which have already been fully assembled and matured.

### ANA1 is important for maintenance of centriole structural integrity downstream of the PCM

If ANA1 is capable of providing stability to centrioles without the maintenance/recruitment of PCM, then it is possible that ANA1 is downstream of the PCM. The PCM may recruit or stabilize components within the centriole structure. If that is the case, we would expect that overexpression of ANA1 should at least partially rescue the loss of centrioles induced by PCM depletion. To test this hypothesis, we used the “centriole stability assay” in culture cells as previously, and overexpressed ANA1 in PCM depleted cells **(Fig. 3 A)**. Interestingly, overexpressing ANA1-GFP in “All PCM” depleted cells led to a reduction of the percentage of cells with abnormally low numbers of centrioles (0-1), evaluated by both BLD10 **(Fig. 5)** and SAS4 markers **(Fig. 5 A, and Fig S 3)**. Indeed, this rescue was total for the BLD10 marker, which reinforces our observations that ANA1 is maintaining the centriole structure in the germline. Additionally, our observations suggest that as it was shown for the PCM, ANA1 is also critical for maintaining the homeostasis of centriolar components (BLD10 and SAS4; **Fig. 5 A and Fig. S 3**). Importantly, simultaneous depletion of PCM (“All PCM” RNAi) and ANA1 has a significantly stronger effect on the maintenance of centriole integrity in comparison with their individual depletions (**Fig. 3 B** and **Fig. S 2 A, B)**. Altogether, our data suggests that ANA1 is important for maintaining the integrity of the centriole structure and that the PCM reinforces that role, perhaps through facilitating the incorporation of newly transcribed ANA1 at the centriole.

**Figure 5-.**
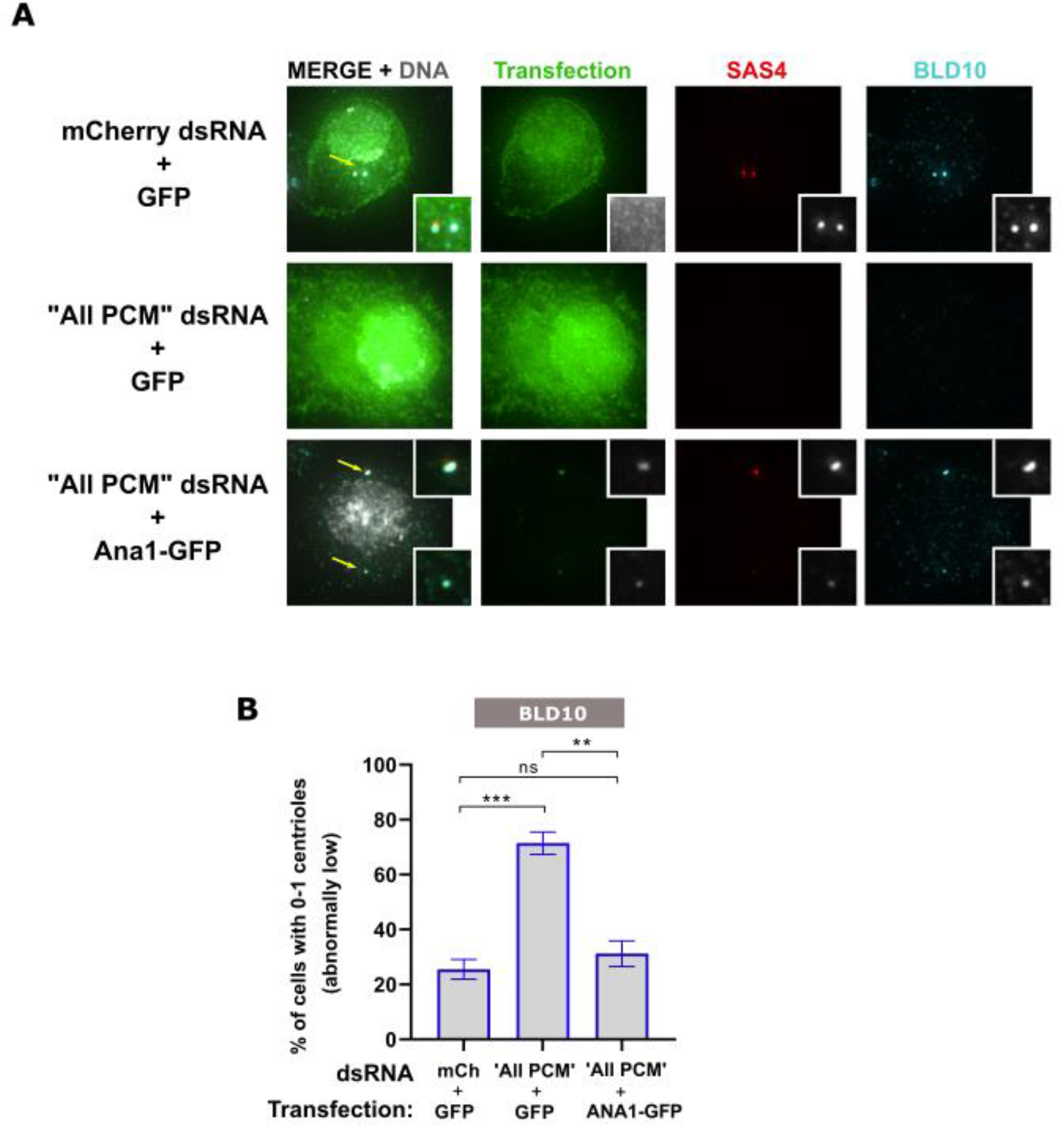
ANA1 rescues the loss of centrioles induced by PCM depletion. **(A)** Cells were stained for SAS4 (red), BLD10 (cyan) and DNA (grey). Representative images are shown. Enlargements of the indicated areas are shown. Scale bar, 10 µm. **(B)** Histograms show the percentage of cells with abnormally low numbers (i.e. 0-1). Centrioles were identified by considering the positive staining in each cell for the centriolar wall protein BLD10. Bars represent the mean ± SEM of three independent experiments (n>40 cells per condition in each experiment). One-way ANOVA, with Tukey’s multiple comparisons test (*, *p*<0.05; ** *p*< 0.01; *** *p*<0.001; ns, not statistically significant). Note that ANA1-GFP rescues centriole loss in the context of PCM depletion.

## Conclusion

Our work suggests that the centriole structure is more dynamic than previously thought and that the turnover of critical components is needed to maintain the integrity of the centriole structure. Such components include proteins from the centriole wall and the cartwheel, as well as the PCM that surrounds the centriole, and Polo kinase that recruits the PCM.

Amongst those components, we show that the conserved centriolar wall protein ANA1 plays a critical role in the maintenance of fully matured centrioles in cultured cells and in the female germline. We further show that Polo and the PCM are dependent on ANA1 for their function in centrosome maintenance: (1) in both germ-line and tissue culture cells, ANA1 RNAi depletion impairs Polo-induced centrosome maintenance, reducing both PCM and centriole protein levels; (2) tethering of ANA1 to the oocyte centrioles is sufficient for their maintenance up to stages where they are normally absent, even in the absence of additional amounts of PCM; (3) overexpression of ANA1 in S-phase arrested culture cells depleted of “All PCM”, rescues the centriole loss normally observed in that condition, showing that ANA1 is a critical component of the PCM-promoted centriole integrity pathway. Our work shows that ANA1 is a critical component for centriole structural integrity, whereas Polo and the PCM are critical components for the maintenance of both centriole structure and centrosome function.

Here we show that ANA1 is not just necessary for centriole biogenesis (Blachon *et al*., 2008, 2009; Dobbelaere *et al*., 2008) and for centriole-to-centrosome conversion (Fu *et al*., 2016), it is also an essential player in the maintenance of centriole integrity and therefore, a critical protein for the entire lifespan of the centrosome. Very recently, the use of several superresolution techniques revealed that ANA1 localises from the pinheads to the outer edge of the doublet microtubules (Tian *et al*., 2021). ANA1 contains multiple predicted coiled-coil regions (Saurya *et al*., 2016) which could potentially promote the interaction with different centrosomal players. One possibility is that ANA1 maintains the microtubule doublets “linked” to each other, promoting the integrity of the wall. This can possibly occur through direct binding of ANA1 to the microtubule doublets (Chang *et al*., 2016), indirectly by its interaction with proteins which provide such links such as BLD10 (Carvalho-Santos *et al*., 2012), or through both pathways. Additionally, ANA1 may also promote cartwheel stability as it was found to interact with SAS6 and ANA2 by yeast two-hybrid (Galletta *et al*., 2016). This central role would also explain the observations made in our candidate screen, where depletion of ANA1 had one of the strongest effects on centrosome maintenance.

How can Polo and the PCM contribute to centriole integrity? *In vitro* work in *C*.*elegans* showed that the PCM can function as a condensate with self-assembling properties, allowing the selective concentration of different components. This was enhanced by Polo/PLK1(Woodruff *et al*., 2017). Moreover, in Drosophila egg extracts, γ-tubulin was proposed to concentrate components to promote *de novo* centriole biogenesis (Nabais *et al*., 2021). Given that ANA1 overexpression rescues the centriole instability phenotype resulting from PCM loss, we hypothesise that the PCM allows a stable concentration of centriolar proteins, and/or regulates their turnover rate with a cytoplasmic pool, which is required for the maintenance of centriole integrity. How can Polo contribute to this pathway? ANA1 phosphorylation at predicted Polo-box binding domains was recently shown to be required for Polo recruitment at the centriole (Alvarez-Rodrigo *et al*., 2021). However, centrioles from flies in which ANA1-dependent Polo recruitment is impaired (mutants expressing an unphosphorylated form of ANA1) are indistinguishable from WT centrioles by EM (Alvarez-Rodrigo *et al*., 2021). Moreover, these flies rescued the uncoordinated phenotype of ANA1 mutants, suggesting that the centriole at the basal body is assembled and properly maintained (Alvarez-Rodrigo *et al*., 2021). Therefore, this phosphorylation at predicted ANA1 Polo-box binding domains does not seem to be critical for centriole integrity. We thus propose that Polo functions in centriole integrity through its role in PCM maintenance.

Oocytes lose their centrioles before fertilisation. Non cycling cells, such as neurons, muscle, and epithelial cells, often attenuate the activity of the centrosome as a MTOC via loss of PCM (Muroyama and Lechler, 2017; Sanchez and Feldman, 2017; Tillery *et al*., 2018). In these cells it is not clear whether the centrioles are eventually lost or remain inactive throughout their whole lifespan. Moreover, it has been suggested that centrosome inactivation is a mechanism used by cancer cells to silence supernumerary centrosomes that otherwise could lead to cell death (Sabino *et al*., 2015). Our work shows that while misregulation of Polo in eggs can lead to infertility in flies, that is not the case for ANA1, as ectopic expression of ANA1 is not sufficient for retaining centrosome function. In future studies it will be of critical importance to address how deregulation of proteins such as ANA1, Polo and the PCM might differentially change the integrity/activity of centrosomes in these contexts and possibly contribute to disease, in particular infertility and cancer.

## Experimental Procedures

### *Drosophila* stocks and genetics Fly stocks

The following fly stocks were used in this study: from Bloomington Stock Centre: UASp-DegradFP (y^1^ w*; M{UASp-Nslmb.vhhGFP4}ZH-51D; #58740); Df(3R)Exel7357 (W^1118^; Df(3R)Exel7357/TM6B, Tb^1^; a deficiency for the *ANA1* locus #7948); Ubi-GFP (y^1^ w^67c23^ P{Ubi-GFP.D}ID-1; #1681); UAS-ANA1-RNAi (y^1^ v^1^ ;P{TRiP.HMJ23356}attP40; #61867, UAS-mCherry-RNAi (y^1^ v^1^ sc* sev^21^;; P{VALIUM20-mCherry}attP2 #35785) and matalpha4-Gal4 (w*; P{matalpha4-GAL-VP16}V2H; #7062). Furthermore, W; ANA1-GFP/CyO and W; BLD10-GFP/CyO (Blachon *et al*., 2008); *ana1*^*mecB*^ mutant flies (Blachon *et al*., 2008); the maternal germline-specific G302-Gal4/TM6B kindly provided by Daniel St. Johnston; Gordon Institute, UK); w; UASp-GFP-PACT; w; UASp-GFP-Polo-PACT and W;;UASp-GFP-Polo.

The following trans-genes were generated for this study:

### Nanobody construct

The coding sequences of PACT (Gillingham and Munro, 2000; Pimenta-Marques *et al*., 2016) and vhhGFP4 ((pGEX6P1-GFP-Nanobody was kindly provided by Kazuhisa Nakayama (Addgene plasmid # 61838)) were PCR amplified (primers used are provided on supplementary table 2). PCR products were purified, excised independently, ligated into the pSpark®-TA Done vector (Canvax), and then transformed into Escherichia coli ‘DH5α’ competent cells. Inserts from at least two different clones were sequenced by the Sanger method. The generated construct (pSpark_GFPnanobody-PACT) was linearized with the restriction enzyme KpnI and cloned into the pDONR™ 221 Vector (ThermoFisher #12536017) Using the gateway® system to generate an entry clone. To create the expression vectors, recombination reactions were achieved using the created entry vector and the destination vector pUbq-phi31. Transgenic flies were generated via plasmid injection (BestGene, INC) using the pUbq-phi31_GFPnanobody-PACT.

The following combination of fly genotypes were generated for this study: *ana1*^*mecB*^ was recombined to G302-Gal4, generating flies W; ; ana1^mecB^ G302-Gal4 Ana1-GFP; ana1^mecB^ G302-Gal4 UASp-DegradFP; Df(3R)Exel7357 Matalpha4-Gal4; UAS-mCherry-RNAi UAS-ANA1-RNAi; UASp-GFP-PACT UAS-ANA1-RNAi; UASp-GFP-Polo-PACT Ubi-GFPnanobodyPACT; G302-Gal4

### Fly husbandry

To degrade ANA1-GFP specifically in the female germ-line at stages 3-4 of oogenesis, flies of the genotype Ana1-GFP; ana1^mecB^ G302-Gal4 were crossed to flies of the genotype UASp-DegradFP; Df(3R)Exel7357. As control, flies of the genotype Ana1-GFP; ana1^mecB^ G302-Gal4 were crossed to Df(3R)Exel7357.

To deplete ANA1 specifically in the female germ-line in the context of Polo-mediated forced maintenance of centrioles, the following crosses were performed: Flies of the genotypes UAS-ANA1-RNAi; UASp-GFP-Polo-PACT and UAS-ANA1-RNAi; UASp-GFP-PACT were crossed to UAS-Matalpha4-Gal4. As controls, Matalpha4-Gal4; UAS-mCherry-RNAi were crossed to either UASp-GFP-Polo-PACT or UASp-GFP-PACT.

To tether different GFP-tagged proteins to the oocytes centrioles in the female germ-line, flies of the genotype Ubi-GFPnanobodyPACT; G302-Gal4 were crossed to flies of the following genotypes: ANA1-GFP/CyO; UASp-GFP-Polo; BLD10-GFP and Ubi-GFP as control.

All strains were raised on standard medium at 25°C, using standard techniques.

### Egg laying and hatching

Single well-fed virgin females with 1 day old were mated with two w^1118^ males in cages with agar plates supplemented with apple juice. The number of eggs laid was counted for 6 days. Each plate was kept at 25°C for 3 extra days and examined for the number of larvae hatched. Egg hatching rates were calculated as the percentage of larvae hatched from the total number of eggs laid by each female. More than 4 independent crosses were performed for each phenotype.

### Ovaries immunostaining

Ovary stainings were performed as previously described (Pimenta-Marques *et al*., 2016). Briefly, females were transferred to pre-warmed (25°C) BRB80 buffer (80 mM Pipes pH 6.8, 1 mM MgCl2, 1 mM EGTA) supplemented with 1X protease inhibitors (Roche), and their ovaries were extracted with pre-cleaned forceps. Individualized ovaries were then incubated for 1 hour (h) at 25°C in BRB80 with 1% Triton X-100 without agitation, followed by a 15 minutes (min) fixation step at - 20°C in chilled methanol. 3 wash steps of 15 min each and overnight permeabilization were done in PBST (1X PBS with 0.1% Tween). Blocking for 1 h was done in PBST with 2% BSA (Gibco). Primary antibodies were incubated overnight at 4°C in PBS with 1% BSA (PBSB) followed by 3 wash steps. Secondary antibodies were diluted in PBSB and incubated for 2 h at room temperature (RT). Ovaries were washed in PBS and DNA was counterstained with DAPI.

### Imaging, analysis and quantification

Drosophila egg chambers were imaged as Z-series (0.3 µm z-interval) on a Zeiss LSM 980, using confocal mode. All images were acquired with the same exposure. Images were processed as sum-intensity projections and intensity measurements were performed using ImageJ software (NIH). Centrosomal regions were determined by the colocalization of at least two different centriolar markers and the intensity of the different proteins were analysed in these colocalization dots. To assess the background level, the intensity of 3 different regions was measured and subtracted to the centrosomal region. In experiments where presence/absence of signal was evaluated, presence of signal was defined as a significant signal above the oocyte background. Image panels were assembled using QuickFigures (Mazo, 2021).

### Protein depletion in DMEL cells

*Drosophila melanogaster* culture cells (DMEL) were maintained in Express5 SFM medium (Gibco, UK), supplemented with 2mM L-Glutamine (ThermoFisher Scientific, UK). Double-stranded RNA (dsRNA) were performed as previously described (Bettencourt-Dias *et al*., 2004). 10 million cells were used for dsRNA transient transfection. dsRNA amounts used in the screen: 20 µg individual *CNN, ASL, D-PLP*, and *SPD2* for “All PCM”; 40 µg *PLK4*, 40 µg *CP110*, 40 µg *CEP97*,40 µg *ANA2*, 40 µg *SAS6*, 40 µg *ANA1*, 40 µg *BLD10*, 40 µg *SAS4*, 80 µg *mCherry* dsRNA. dsRNA combination amounts used for Figure 3: “All PCM” combined with 20 µg of *mCherry* dsRNA or 20 µg of *ANA1* 5’-3’-UTR dsRNA; 100 µg of *mCherry* dsRNA. Primers used for dsRNA production are listed in Supplementary Table 1.

### “Centriole Stability Assay”

Centriole stability assay was performed as previously described (Pimenta-Marques *et al*., 2016) (Pimenta-Marques et al., 2016). Briefly, DMEL cells were S-phase-arrested with 10 µM aphidicolin (Aph), a specific eukaryotic DNA polymerase inhibitor, and 1.5 mM hydroxyurea (HU), which reduces deoxyribonucleotide production, 1h after dsRNA transfection. For 8 days “centriole stability assay” (Fig. 1), cells were subject to a second round of dsRNA transfection and Aph+HU treatment after 4 days. For 4 days “centriole stability assay” (Fig. 3 and Fig. 5), cells were collected after 4 days of dsRNA and Aph + HU treatment.

### Plasmid transfections

DMEL cells were transiently transfected with GFP-PACT, GFP-POLO-WT-PACT, GFP, or ANA1-GFP after transfection with dsRNA either *mCherry* dsRNA (control), *“All PCM”* dsRNA or “*All PCM” + ANA1* dsRNA. Since GFP-PACT and GFP-POLO-WT-PACT constructs contain an UASp promoter, each of these constructs were simultaneously co-transfected with an Actin5C-Gal4 plasmid. Plasmid transfections were performed as previously described (Pimenta-Marques *et al*., 2016).

### Immunostaining and imaging of *D. melanogaster* culture cells

DMEL cells were plated into glass coverslips and allowed to adhere for 1h at 25°C. Cells were then fixed for 10 min at room temperature with a solution containing 4% paraformaldehyde, 60 mM PIPES pH 6.8, 30 mM HEPES pH7.0, 10 mM EGTA pH 6.8, 4mM MgSO_4_. After 3 washes, cells were permeabilized and blocked with PBSTB (a PBS solution containing 0.1% Triton X-100 and 1% BSA). Cells were incubated with primary antibodies diluted in PBSTB overnight at 4°C. After 3 washes, cells were incubated with secondary antibodies and DAPI (Santa Cruz Biotechnology) diluted in PBSTB for 2H at RT. Cells were mounted with Vectashield Mounting Medium (Vector laboratories). Cell imaging was performed in Deltavision OMX (Deltavision) microscope with a PCO Edge 5.5 sCMOS 2560×2160 camera, or with a Nikon High Content Screening (Nikon) microscope with an Andor Zyla 4.2 sCMOS 4.2Mpx. All images were acquired with the same exposure. Images were acquired as Z-series (0.2 µm z-interval) and analysed as maximum intensity projections. Image panels were assembled using QuickFigures (Mazo, 2021).

### Antibodies

Primary antibodies and dilutions used: chicken anti-PLP (1:1000 for DMEL cells immunostaining; 1:500 for ovary immunostaining), kindly provided by David Glover,University of Cambridge, UK (Bettencourt-Dias *et al*., 2005); rabbit anti-Bld10 (1:2000), kindly provided by Timothy Megraw, The Florida State University, USA (Mottier-Pavie and Megraw, 2009); rat anti-ANA1 (1:500), kindly provided by Jordan Raff, University of Oxford, UK (Saurya *et al*., 2016); mouse anti-γ-tubulin (1:50 dilution; clone GTU88, Sigma); rabbit anti-SAS4 (1:500, Metabion); rabbit anti-CP110 (1:10000 for DMEL cells immunostaining; 1:5000 for ovary immunostaining, Metabion). Secondary antibodies (Jackson Immunoresearch Europe) were used at 1:1000 for Dmel cells immunostaining, and 1:250 for ovaries immunostaining.

## Acknowledgments

We thank all members of the Cell Cycle and Regulation Lab for discussions and for critical reading of the manuscript. We thank Tomer Avidor-Reiss (University of Toledo, Toledo, OH), Daniel St. Johnston (The Gurdon Institute, Cambridge, UK), David Glover (University of Cambridge, Cambridge, UK), Jingyan Fu (Agricultural University, Beijing, China) Jordan Raff (University of Oxford, Oxford, UK) and Timothy Megraw (Florida State University, Tallahassee, FL) for sharing tools. We acknowledge the technical support of Instituto Gulbenkian de Ciência (IGC)’s Advanced Imaging Facility, in particular Gabriel Martins, Nuno Pimpão Martins and José Marques. We also thank Tiago Paixão from the IGC’s Quantitative & Digital Science Unit and Marco Louro from the CCR lab. for the support provided on statistical analysis. IGC’s Advanced Imaging Facility (AIF-UIC) is supported by the national Portuguese funding ref# PPBI-POCI-01-0145-FEDER -022122. We thank the IGC’s Fly Facility, supported by CONGENTO (LISBOA-01-0145-FEDER-022170). This work was supported by an ERC grant (ERC-2015-CoG-683258) awarded to M. Bettencourt-Dias and a grant from the Portuguese Research Council (FCT) awarded to A. Pimenta-Marques (PTDC/BIA-BID/32225/2017).

## Authors Contribution

Conceptualization: A. Pimenta-Marques, T. Perestrelo, Bettencourt-Dias. Methodology: Patrícia Rodrigues and A. Pimenta-Marques (candidate screen in *Drosophila* cultured cells); A. Pimenta-Marques (deGradFP experiments in oogenesis, depletion of ANA1 in GFP-Polo-PACT expressing flies, and GFPnanoPACT experiments in oogenesis) and T. Perestrelo (depletion and rescue experiments in *Drosophila* cultured cells). Validation: A. Pimenta-Marques, T. Perestrelo, P. Rodrigues, P. Duarte, M. Lince-Faria and M. Bettencourt-Dias. Investigation: Patrícia Rodrigues (candidate screen in *Drosophila* cultured cells); A. Pimenta-Marques (depletion of ANA1 in GFP-Polo-PACT expressing flies, and GFPnanoPACT experiments in oogenesis) and T. Perestrelo (deGradFP experiment in oogenesis and depletion and rescue experiments in *Drosophila* cultured cells). Analysis: A. Pimenta-Marques, T. Perestrelo, Patricia Rodrigues. Investigation: Resources: A. Pimenta-Marques (generation of fly genotype combinations used in this study) and T. Perestrelo and P. Duarte (molecular biology - generation of tools used in different experiments). Writing – original draft: A. Pimenta-Marques and T. Perestrelo – Reviewing and editing: A. Pimenta Marques, T. Perestrelo, P. Rodrigues, M. Lince-Faria and M. Bettencourt-Dias. Supervision and coordination A. Pimenta-Marques and M. Bettencourt-Dias.

## Conflict of Interest

The authors declare no conflict of interest.

---

**Figure S1.**
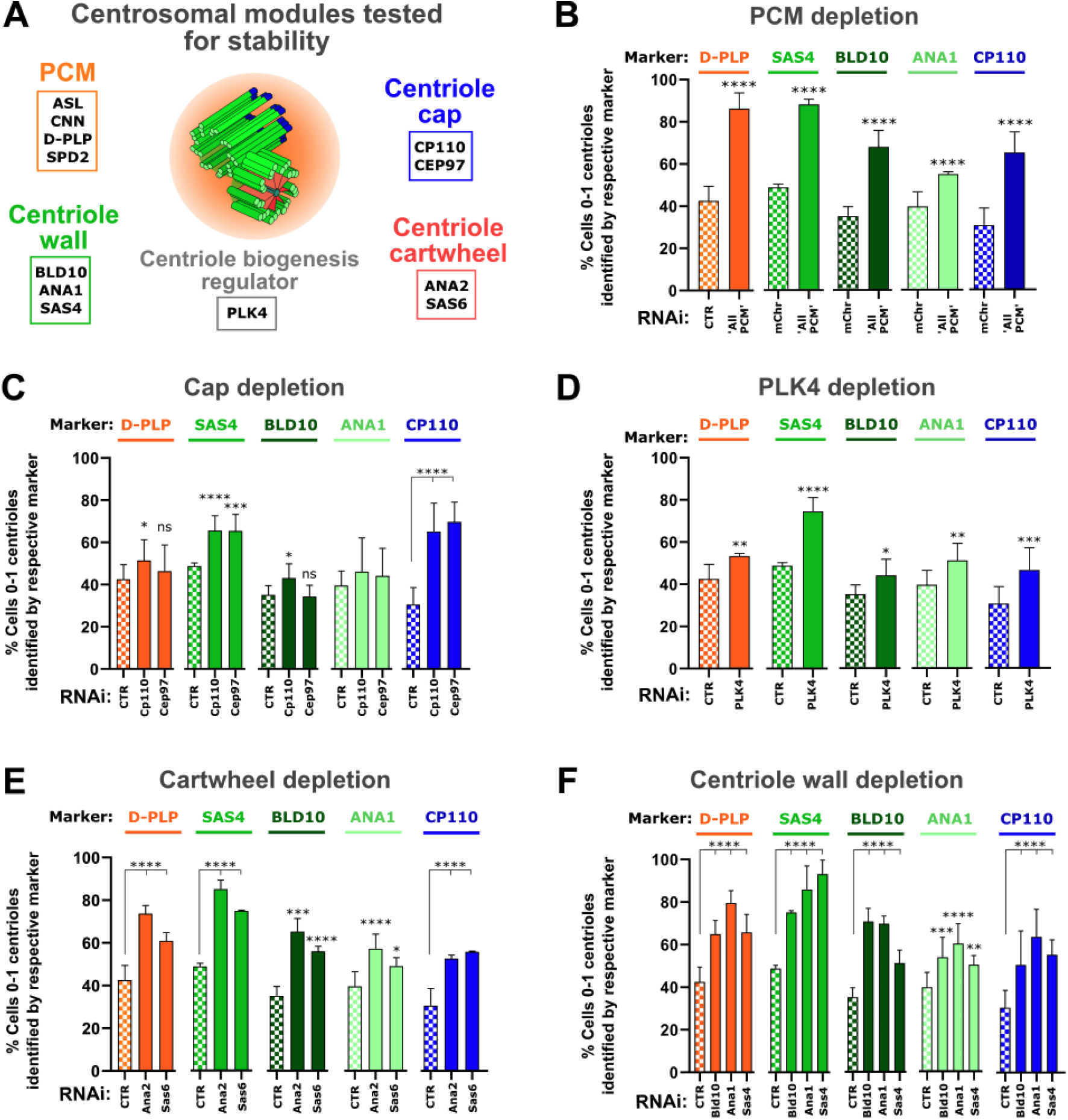
Candidate screen in *Drosophila* cultured cells for centrosome maintenance. **(A)** Schematic representation of the different proteins that were depleted in each of the centrosome modules tested for maintenance. These include: “ALL PCM” proteins (simultaneous depletion of four major PCM proteins: ASL, CNN, D-PLP and SPD-2); centriole cap proteins (CEP97 and CP110); the major regulator of centriole biogenesis, PLK4; cartwheel proteins (ANA2 and SAS6) and the centriolar wall proteins (BLD10, SAS4 and ANA1). **(B-F)** Centriolar numbers were assessed considering the positive staining in each cell for different centrosome markers. These include: the PCM marker D-PLP (orange bars); the centriole wall markers SAS4 (green bars), BLD10 (dark green bars) and ANA1 (light green bars) and the distal cap protein CP110 marker (blue bars). Histograms represent the percentage of cells with abnormally low centriole numbers (i.e. 0-1) **(B)** Depletion of “All PCM”, **(C)** Depletion of the centriolar cap proteins CP110 or CEP97; **(D)** Depletion of the centriolar biogenesis regulator PLK4, **(E)** Depletion of the cartwheel proteins ANA2 or SAS6, and **(F)** depletion of the centriolar wall proteins BLD10, ANA1, or SAS4. Bars represent the mean ± SEM of three independent experiments (n ≥ 100 cells per condition in each experiment). Generalised linear binomial model in which the effect of each replicate and RNAi were considered as independent factors. * *p*< 0.05; ** *p* < 0.01; \*\*\**p* < 0.001; \*\*\*\**p* < 0.0001; ns, not significant (see also **Fig. 1**).

**Figure S2.**
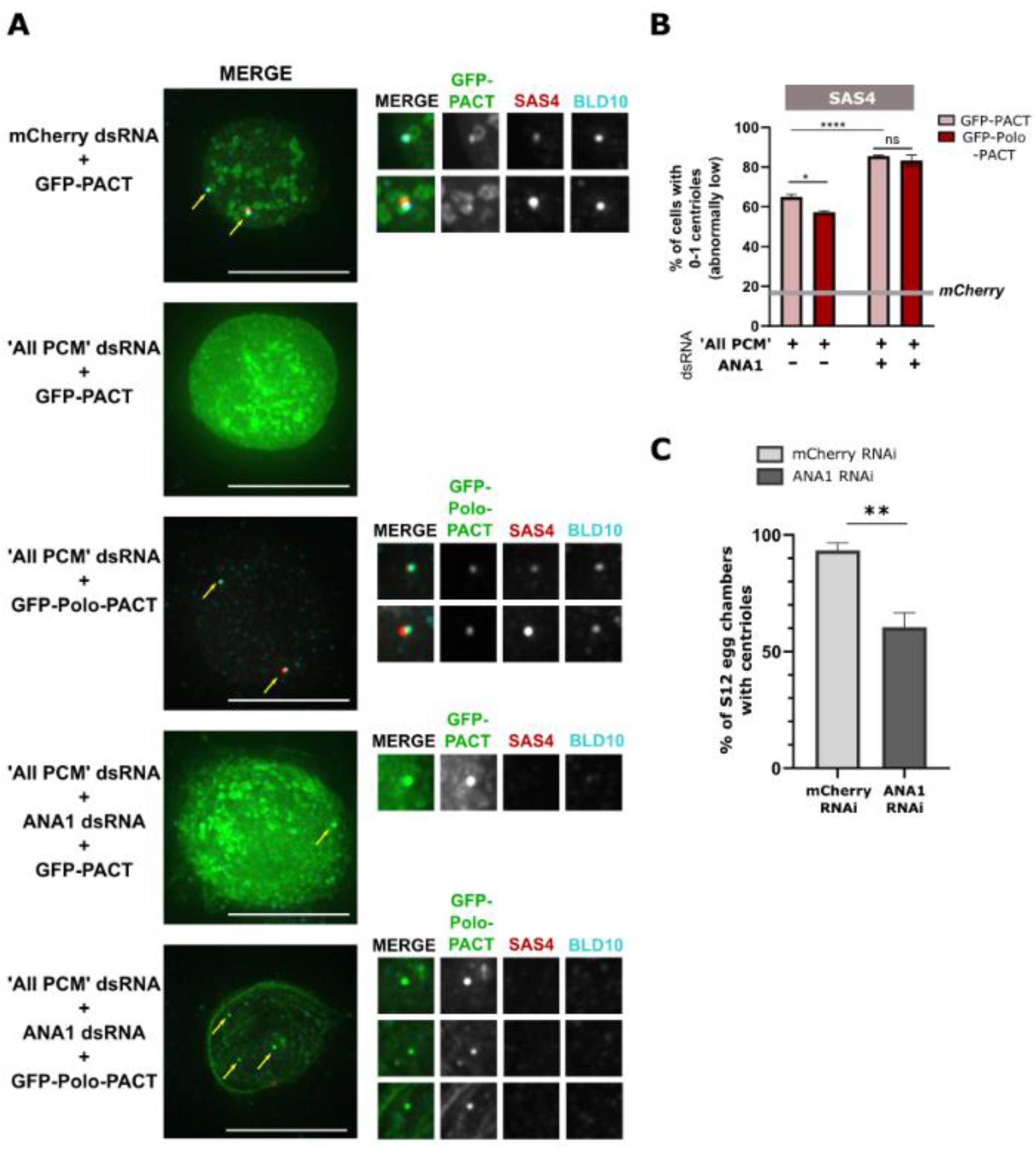
Polo kinase induced centriole stability is dependent on ANA1. **(A)** DMEL cells arrested in S-phase (treatment with hydroxyurea (HU) and aphidicolin (Aph) were depleted of “All PCM” (simultaneous depletion of ASL, CNN, D-PLP and SPDS-2), or simultaneous depletion of “All PCM” and ANA1, and transfected with either GFP-PACT or GFP-Polo-PACT. mCherry dsRNAwas used as a negative control. Cells were immunostained for BLD10 (cyan), SAS4 (red). Representative images are shown. All conditions were acquired with the same exposure. Enlargements of the indicated areas are shown. Note that although GFP aggregates are present when “All PCM” and ANA1 are co-depleted, they do not correspond to centrioles as there is no colocalization between GFP aggregates and BLD10 and/or SAS4. Scale bar, 10 µm. **(B)** Histograms show the percentage of cells with abnormally low numbers (i.e. 0-1) of the indicated centriole marker SAS4. Bars represent the mean ± SEM of three independent experiments (n>100 cells per condition in each experiment). Two-way ANOVA, with Tukey’s multiple comparisons test (*, *p*<0.05; \*\*\*\**p*<0.0001; ns, not statistically significant). **(C)** Stage 12 egg chambers expressing GFP-Polo-PACT (centrosome maintenance conditions) with *mCherry* RNAi (control for RNAi, n=28) or ANA1 RNAi (n=27). Statistical significance was determined by Fisher’s Exact test on the pooled replicates, after testing whether there were significant differences between them. Shown are mean ± SEM. (see also **Fig. 3**).

**Figure S3.**
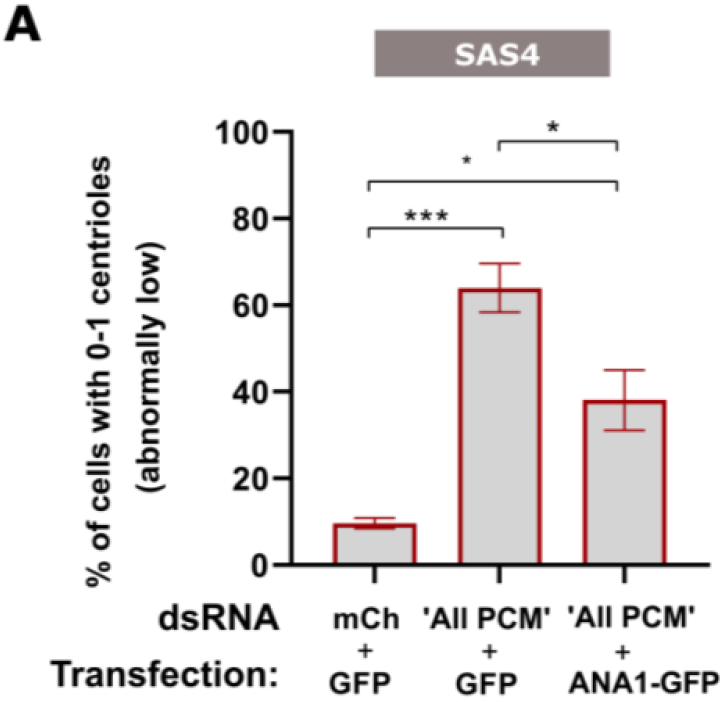
ANA1 rescues the loss of centriole maintenance induced by PCM depletion. **(A)** Histograms show the percentage of cells with abnormally low numbers (i.e. 0-1). Centrioles were identified by considering the positive staining in each cell for the centriolar wall protein SAS4. Bars represent the mean ± SEM of three independent experiments (n>40 cells per condition in each experiment). One-way ANOVA, with Tukey’s multiple comparisons test (*, *p*<0.05; ** *p*< 0.01; *** *p*<0.001; ns, not statistically significant). Note that expression of ANA1-GFP rescues centriole loss in the context of PCM depletion.

**Table 1.**
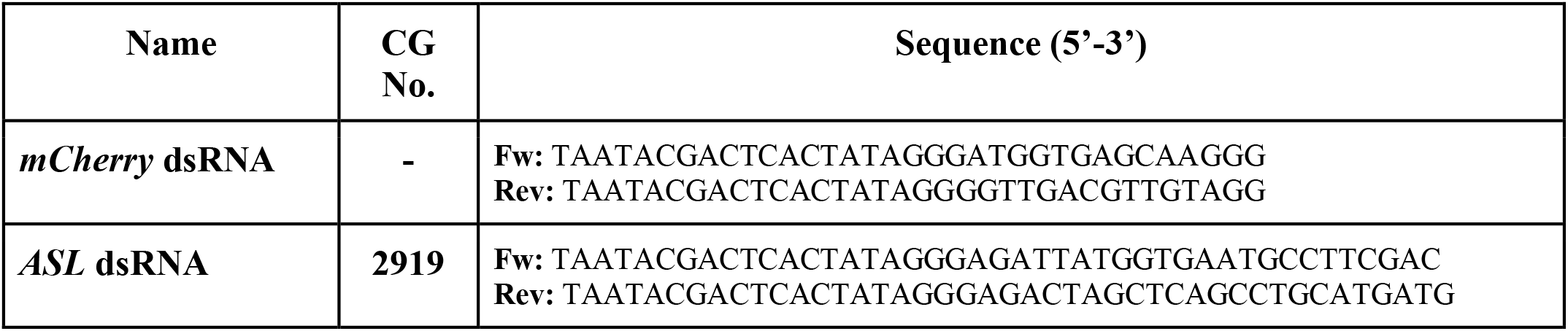
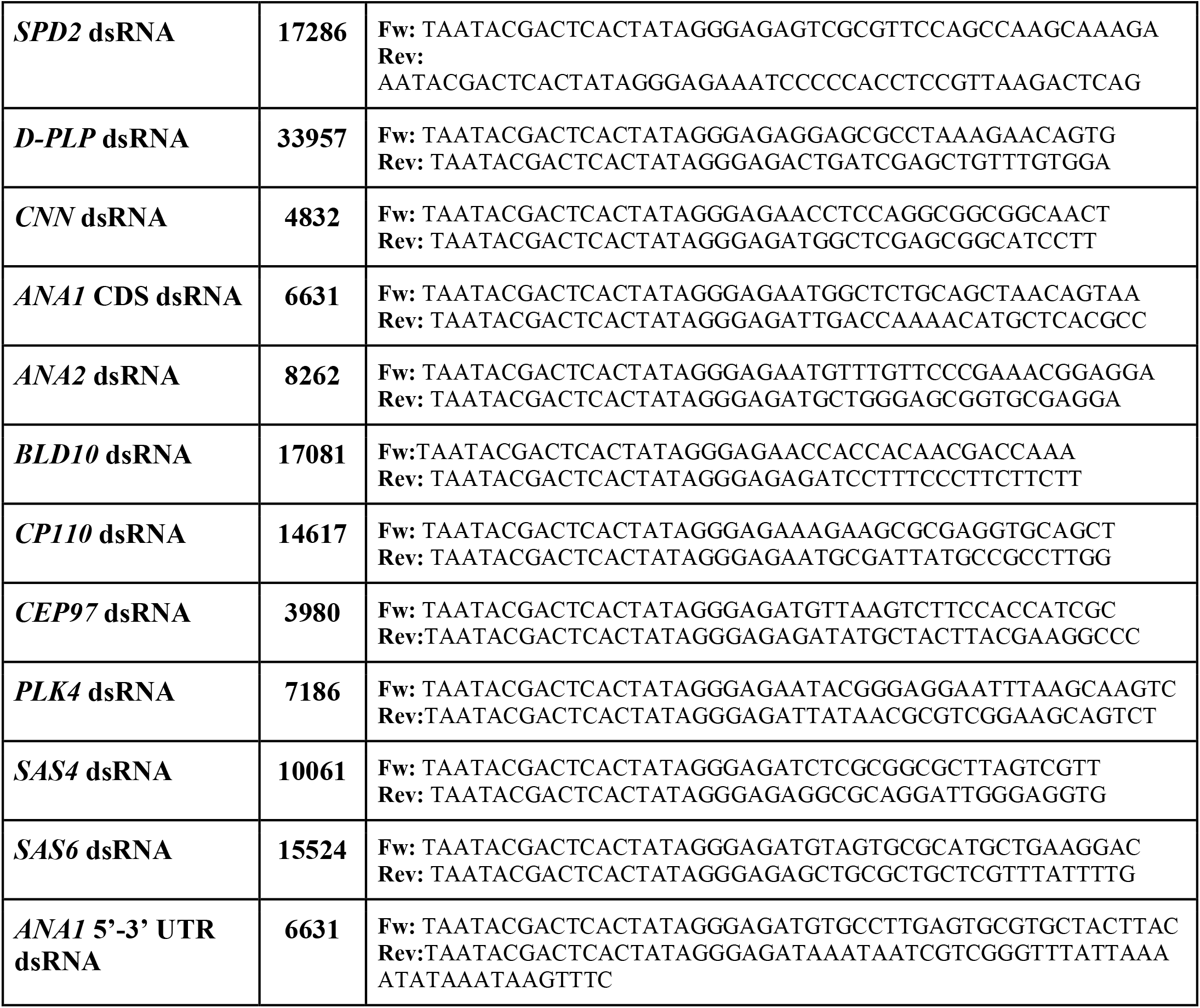
List of primers used for dsRNA synthesis.

## Full sequence of 5′-3′-UTR-3′UTR hybrid as template for dsRNA

**TAATACGACTCACTATAGGGAGA**TGTGCCTTGAGTGCGTGCTACTTACCAGCTGGTATATTTTAGACGCATGTAAATTCTAGTACATTCAATTATTCATCTACGGTCACACTGCCGCTTGGGAGGAATTTTAAAGACGTTGGGTTGTTTGATTTTACGCTCAAACTTGTTTCGATTTCTACTGCGTAAATGCTGCCCCACATACGAATTTATTACATATATCGATAGAGCAGTCGCCGAACTTTTAATTCGTTTTTTAGGTTTTAGATTATATTATCCATTTTATGACAATTATTTATATTTTACTTACTTTGCAATTTTGTGTCAAAAAATGACTATCGAAAAAGATTGTATAAAATTTACTCAATAAGTTAAATGTACATTTTATTACCAATTTGTGTGAAACTTATTTATATTTTAATAAACCCGACGATTATTTAT**TCTCCCTATAGTGAGTCGTATTA**

**Table 2.**
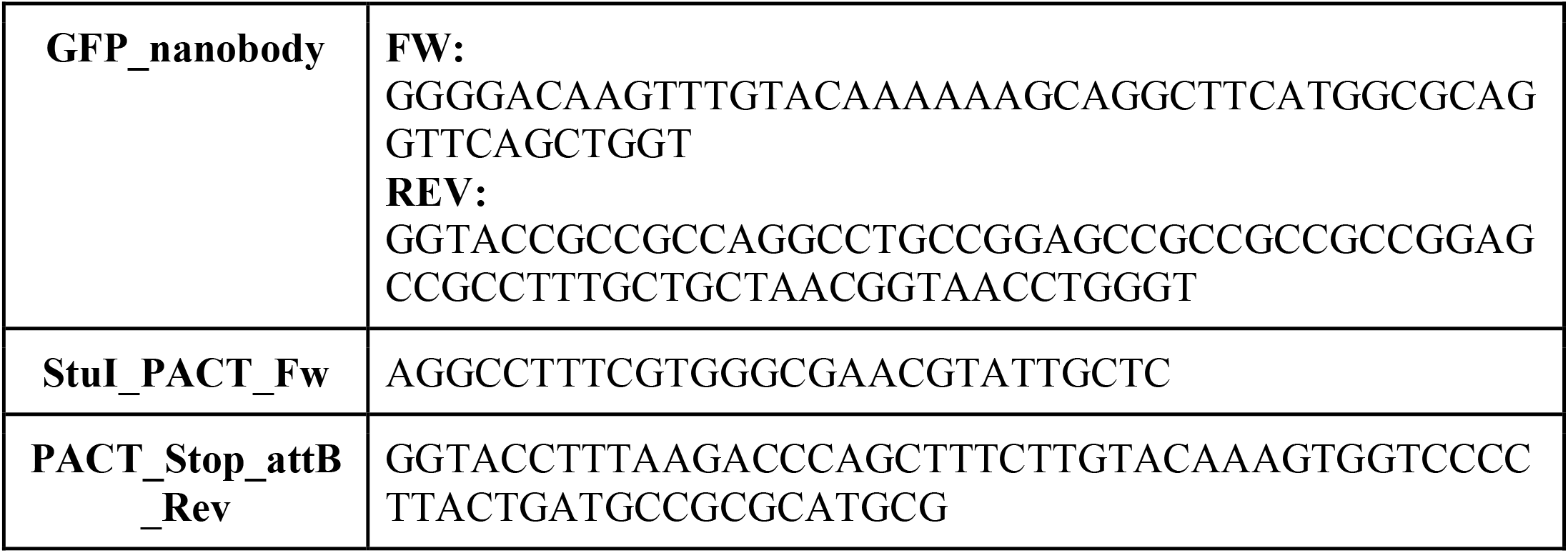
List of primers used for generation of GFP-nanoPACT construct.

## Notes

### Competing Interest Statement

The authors have declared no competing interest.

